# Distinct sub-MIC kill kinetics of Cu and Ag in *Escherichia coli*

**DOI:** 10.1101/2025.08.27.672559

**Authors:** Merilin Rosenberg, Sigrit Umerov, Carmen Marianne Teär, Angela Ivask

## Abstract

Copper and silver are well-known and widely used antimicrobial metals that are often considered to employ diverse overlapping biocidal mechanisms of action and induce corresponding bacterial defense responses. Exposure to antimicrobial metals at concentrations below the minimal inhibitory concentration (sub-MIC) are widespread in natural, clinical and built environments and can shape evolutionary trajectories of the affected microbes that could lead to antimicrobial tolerance and/or resistance. By analyzing growth and kill kinetics *of Escherichia coli* under copper or silver exposure we observed that sub-MIC copper concentrations resulted in a lasting dose-dependent slowing of exponential growth with reduced yield while silver seemed to cause dose-dependent growth delay without substantially affecting exponential growth or yield. Time-kill experiments revealed minimal loss of viability in early copper exposure while in case of silver a rapid dose-dependent transient killing followed by normal exponential regrowth of the survivors was observed, underlying the seemingly dose-dependently extended lag phase durations. Distinguishing conditions that select for antimicrobial resistance (sustained growth) versus tolerance (survival without growth) is essential for antimicrobial stewardship as acquiring tolerance is considered a steppingstone towards developing resistance. Our results suggest that short-term survival of the initial killing by silver is sufficient for selective advantage while maintaining energy-intensive enhanced growth in the presence of copper is needed to gain competitive benefit over the general population. The findings highlight new and known challenges in antimicrobial characterization and risk assessment of metal-based formulations by using wide-spread non-kinetic endpoint assays such as MIC.

## Main text

### Silver and copper are widely used in various antimicrobial applications with the intent to kill or limit the spread of potentially pathogenic microbes, yet their mechanisms of action are not fully understood

Cellular targets of the antibacterial activity of copper and silver as well as the corresponding bacterial defenses are often considered overlapping due to similar coordination chemistry and sulfhydryl reactivity, leading to shared efflux and sequestration mechanisms (1–5). *E.g*., the CusCFBA system exports both Cu(I) and Ag(I) and is induced by CusRS (5). Unlike toxic silver, copper is an essential micronutrient tightly regulated by homeostasis but becomes toxic in excess (1, 6). A key difference is Cu(I)/Cu(II) redox activity, which generates reactive oxygen species and causes direct damage such as lipid peroxidation (7). We have previously found that silver and copper also differ in their relative toxicity to bacteria depending on exposure conditions. *E. coli* tolerates higher copper ion concentrations (millimolar range) than silver ion concentrations (micromolar range) in liquid exposure media but survives longer on silver surfaces than on copper surfaces in semi-dry exposure conditions (8, 9). These contrasts suggest distinct antibacterial mechanisms or condition-dependent defense responses.

### Antibacterial activity is typically measured and compared via static endpoints not kinetic assays

Minimal inhibitory concentration (MIC) indicates presence or absence of growth after standardized exposure while minimal biocidal concentration (MBC) only reflects endpoint mortality (>99.9%) (10, 11). However, endpoint assays can mask regrowth after transient killing. To capture dynamics, where an antimicrobial agent is quickly detoxified and/or causes transient killing that allows survivors to regrow before the assay endpoint, growth and kill kinetics assays in well-defined experimental conditions should be used. Implications of the strain, media and exposure conditions selection in the current study are further discussed in Supplementary Note 1.

### Silver and copper appear to affect *E. coli* growth kinetics differently

To study the effects of copper and silver on the growth and viability of *E. coli*, exponential cultures were exposed to 0.0625 to1 times the 48 h MIC value increments of the respective metal salts in MOPS-buffered defined medium (Fig. 1; detailed methods of the growth and time-kill experiments presented can be found in Supplementary Note 2). At comparable sub-MIC concentration ranges, CuSO_4_ increased doubling time and reduced biomass yield of *E. coli* in a dose-dependent manner (Fig. 1A, Supplementary Fig. S1 and S2). In contrast, AgNO_3_ seemingly prolonged lag time with little effect on doubling time or yield (Fig. 1B, Supplementary Fig. S1 and S2).

**Figure 1.**
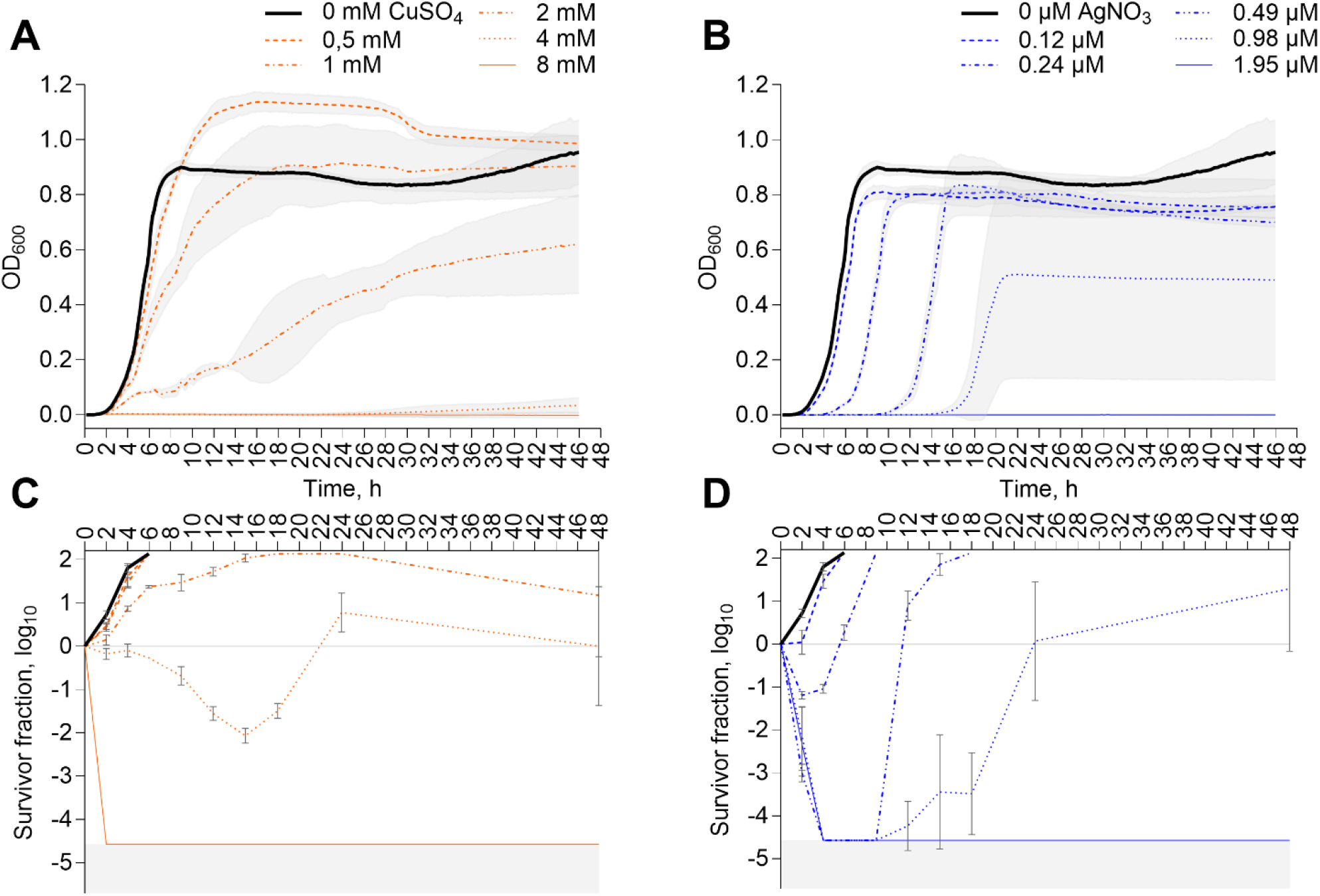
Growth (A, B) and kill (C, D) kinetics of *Escherichia coli* BW25113 exposed to CuSO_4_ (A, C) or AgNO_3_ (B, D) in the range of 0.0625-1 xMIC value of 8 mM Cu or 1.95 µM Ag after 48 h. Mean and SD of 3 (C, D) to 4 (A, B) biological replicates are shown. All growth curves from individual experiments represented on panels A and B can be found in Supplementary Fig S3 and survival data on panels C and D presented as bar graphs on Supplementary Fig S5, correlation between metal concentration and growth parameters in Supplementary Fig S2 and numerical data in Supplementary Data file. Grey shading on A and B denotes standard deviation and on C and D value range below post-exposure colony counting detection limit with no colonies detected.

### Silver and copper cause distinctly different sub-MIC kill kinetics during the ostensible lag phase

Copper exposure resulted in minimal early killing in addition to lasting growth inhibition (Fig. 1A and 1C), whereas silver caused rapid dose-dependent up to over 4-log reduction in viable counts prior to resumption of normal exponential growth (Fig. 1B and 1D). Prolonged physiological lag time can signal a stress response (12, 13) and, without the time-kill data in Fig. 1D, could plausibly account for the delayed growth in Fig. 1B. However, a decrease in inoculum density also increases population-level lag phase duration - a phenomenon commonly known as the inoculum effect (12, 14). The latter was also evident for *E. coli* under our growth-assay conditions (Supplementary Fig. S1 and S4). Therefore, dose-dependently increased lag phase duration under antimicrobial exposures as in case of silver in Fig 1B, could be primarily attributable to sub-MIC killing and reduced viable cell count at resumption of growth as illustrated on Fig. 1D. In case of observed lag shifts, growth kinetics should be complemented by sub-MIC tolerance assays to discriminate partial killing from physiologically prolonged lag phase.

### The results suggest that repeated or prolonged sub-MIC copper and silver exposures could differentially select for tolerance and resistance

Tolerance - survival without growth - is easier to develop than resistance, which requires enhanced growth in the presence of an antimicrobial agent. Tolerance is thus considered a precursor to resistance (15, 16), offering survivors time to acquire adaptive traits. While both silver and copper are used in commercial antimicrobial products (*e.g*. wound care, textiles, touch surface materials) copper resistance is more studied in the context of environmental pollution or colonization of the indoor environment. On the other hand, silver resistance is a well-known challenge necessitating combined treatment strategies, *e.g*. in burn wound care (17, 18). Our results suggest that short-term survival of the initial killing by silver is sufficient to gain competitive benefit over the general population while in the presence of copper overcoming growth inhibition and continuous upkeep of the energy-intensive advantage is needed. This raises the question of whether silver-based antimicrobial applications generally pose a higher risk for the development of tolerance and resistance compared to copper-based applications, due to the distinct growth inhibition and kill kinetics of copper and silver. It also prompts further investigation into how these differences influence the co-selection of antibiotic tolerance and development of antibiotic resistance in medical applications. In conclusion, our observations highlight new and known challenges in assessing antimicrobial characteristics by using wide-spread endpoint assays such as MIC that mask toxicity kinetics to make decisions guiding the design and risk assessment of antimicrobial formulations in different fields of application.

## Acknowledgements

The research is conducted using the research infrastructure “Experimental Studies and Applications of Cellular Processes – RAKERA” funded by the Estonian Research Council (TARISTU24-TK14) and supported by Estonian Research Council grant PRG1496. The study received funding from the European Commission Twinning project FAST-Real (101159721) and the Estonian Ministry of Education and Research under projects TK210 and TEM-TA55.

## Supplementary Notes and Figures for

### Supplementary Note 1 on strain selection, growth and exposure conditions

#### Strain and inoculum selection

We first observed the effects described in this study for *E. coli* ATCC 8739 (unpublished data), a strain that is routinely used in antimicrobial testing. However, the strain tends to rapidly acquire silver resistance in liquid medium granted by mutations in the cryptic genomic *sil* locus (1–4) that interferes with studying the effects of antimicrobial metals on the wild-type strain. *E. coli* K-12 derivative BW25113 (5) was used for the current study as a better described model strain of *E. coli* lacking the *sil* locus. Exponential phase inoculum was used in the study similarly to standardized MIC and MBC assays and to avoid potential biases caused by inoculating stationary or biofilm cultures with different tolerance profiles into fresh growth-supporting exposure medium. Due to the limited growth phase coverage, the results cannot be generalized to stationary and colony-biofilm cultures in which cases additional experiments would be necessary.

#### Exposure medium selection

It has been demonstrated before that transferring *E. coli* from pH=7.0 to pH=4.2 medium can substantially decrease viable counts (6) and pH=5.5 also seems to both decrease growth rate and prolong lag phase at population level compared to pH=7.5 (7). In addition, during exponential growth phase *E. coli* culture itself can further decrease pH of a glucose-supplemented medium by 1.5-2 pH units (8, 9).

In the context of antimicrobial metals, the addition of 365 ppm (5.7 mM) or 650 ppm (10 mM) copper to regular unbuffered medium such as LB has been shown to decreases pH of the medium from 7.2 to 5.3 or 4.7 accordingly (10), an acidity range that impairs growth of *E. coli* also without excess copper. Similarly, also in our experience supplementing LB (Lennox salt) with CuSO_4_ reduces pH while the pH change is expected to be much smaller in the buffered MOPS Minimal Medium (MMM), figure **to the left ←**, whereas silver had no effect on pH of the media (data not shown).

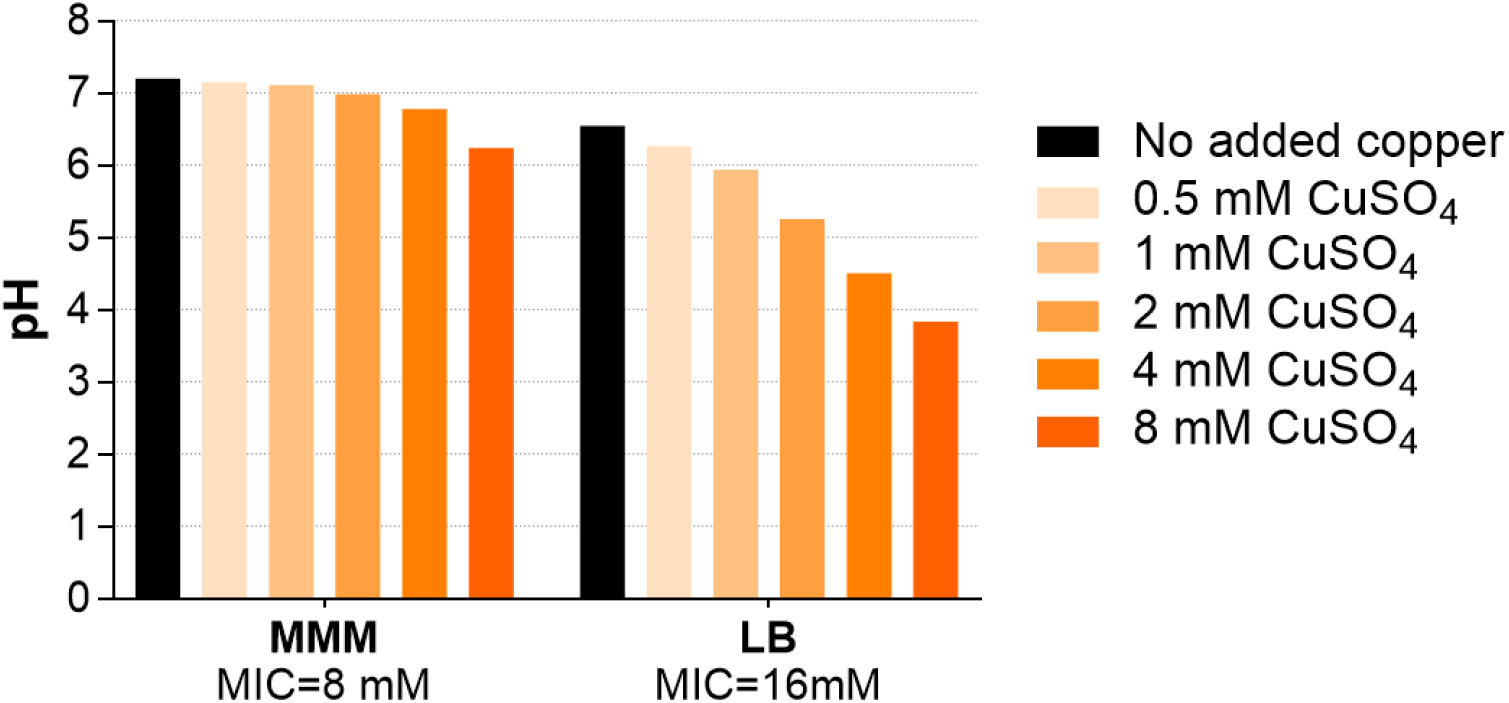

Additionally, acid resistance, pH recovery and growth dynamics of *E. coli* subjected to near-lethal acid stress in unbuffered LB also depends on inoculum density with complete growth inhibition at initial pH≤4.0 and MIC assay cell density while increasing pH to above ∼4.4 is needed before resuming growth (11). This further complicates interpretation of results with varying cell counts or potentially partial initial killing by either acidity or copper or both. Additional toxic effect of acidification due to copper in the medium might spark ideas for combined synergistic use of antimicrobial agents. However, the slow-release copper sources such as CuO nanoparticles do not seem to have a similar effect on medium pH.

MMM (12, 13) was selected for metal exposures due to its buffering capacity to reduce pH bias while comparing antimicrobial activity of copper and silver. Buffered medium counteracts acidification upon addition of CuSO_4_ and helps to maintain the pH near the optimal growth range of *E. coli* (pH ∼6.5-7.5) at sub-MIC copper concentrations. Metal ion toxicity is heavily affected by exposure medium (14, 15). MMM was also used due to its defined characteristics and lower organic content to control the metal ion speciation effects on bioavailability and thus toxicity to bacteria (*e.g*. formation of insoluble salts and complexing of metal ions by thiol groups). Both silver and copper ions can be complexed by organic contents, especially cysteine residues, of the medium with variable affinity, while only silver forms practically insoluble AgCl in the presence of ∼60 mM chloride ions in MMM. However, similar situations would be encountered in clinically relevant conditions. Chloride content of MMM is lower than in LB (Miller 10 g/L NaCl → 171 mM; Lennox 5g /L NaCl → 86 mM), PBS (140 mM) or human blood plasma (∼100 mM).

#### Re-growth conditions

We have previously observed that copper exposure can cause slower colony growth substantially affecting results of agar MIC assays (2) possibly also due to copper-induced pH changes. Therefore, growth and kill kinetics were recorded for 48 h instead of 24 h. Drop-plated colonies from the kill curve experiments were counted after 14-16 h for optimal countable colony size and re-checked after 24, 48 and 72 h. Spread-plates from kill curves were counted after 24 h and checked after 48 and 72 h for possible slower growth from exposures with higher metal concentration. No delayed emergence of colonies was observed. 96-well exposure plates from the growth and kill kinetics experiments were kept at 37°C and 150 rpm for 5 days after the 48-h data collection and visually confirmed that delayed growth (emergence of visible turbidity) was not observed for metal concentrations at which no OD increase had been detected during 48 h.

### Supplementary Note 2 on method details

Escherichia coli K-12 derivative BW25113 was used for the current study as a well-described model strain of *E. coli*. Lysogeny broth (LB: 5 g/L yeast extract, 10 g/L tryptone, 5 g/L NaCl with optional 15 g/L bacteriological agar) and 37°C incubation with or without 150 rpm orbital shaking was used for all precultures and phosphate-buffered saline (PBS: 8 g/L NaCl, 0.2 g/L KCl, 1.44 g/L Na_2_HPO_4_, 0.2 g/L KH_2_PO_4_; pH 7.1) for washing and serial dilutions of cultures. Metal exposures used MOPS Minimal Medium (MMM)(12, 13) supplemented with 0.4% glucose, 0.4% casamino acids, and 20 µg/L tryptophan (∼60 mM chloride in 1× MMM as described above).

Second subculture of *E. coli* on LB was used to inoculate overnight liquid LB cultures. The stationary culture was 100-fold diluted into fresh LB and grown to exponential phase (OD_600_=0.5-0.7), then washed twice with cold PBS (4°C, 4400 g, 10 min). Pellets were resuspended in 2× MMM and adjusted to OD_600_ = 0.002 (∼10^6^ CFU/mL). Inoculum was added to twofold dilution series of AgNO_3_ or CuSO_4_ or mixed 1:1 with deionized water and serially diluted in 1× MMM to test inoculum size effects. The final volume of exposures in 96-well suspension culture plates was 150 µl for growth curves and 200 µl for kill curves. Sterile water was pipetted between the wells and plates sealed with parafilm to reduce drying during long incubations.

Growth curves were measured at OD_600_ every 15 min at 37°C with double-orbital mixing (BioTek Synergy H1, Agilent). Blank-corrected data were analyzed with Dashing Growth Curves at https://dashing-growth-curves.ethz.ch/ (16).

Minimal inhibitory concentrations (MIC) of AgNO_3_ or CuSO_4_ were determined from growth curves as the lowest concentration at which no time-dependent OD increase was registered during the experiments and verified by no visible growth after 48 h incubation.

For kill curves, exposure plates were incubated at 37°C, 150 rpm in the dark. At each time-point, three concentration series per metal were serially diluted in PBS and drop-plated on LB; 150 µL of undiluted exposures were also spread-plated on 10 cm LB plates for viable counts. Survival was expressed as log_10_-transformed survivor fraction (post-/pre-exposure viable counts).

Statistical analysis was done in GraphPad Prism 10.4.1 using correlation, linear regression and one-way ANOVA with post-hoc testing for multiple comparisons at α = 0.05 across 3–4 biological replicates, as detailed in figure legends.

**Supplementary Figure S1.**
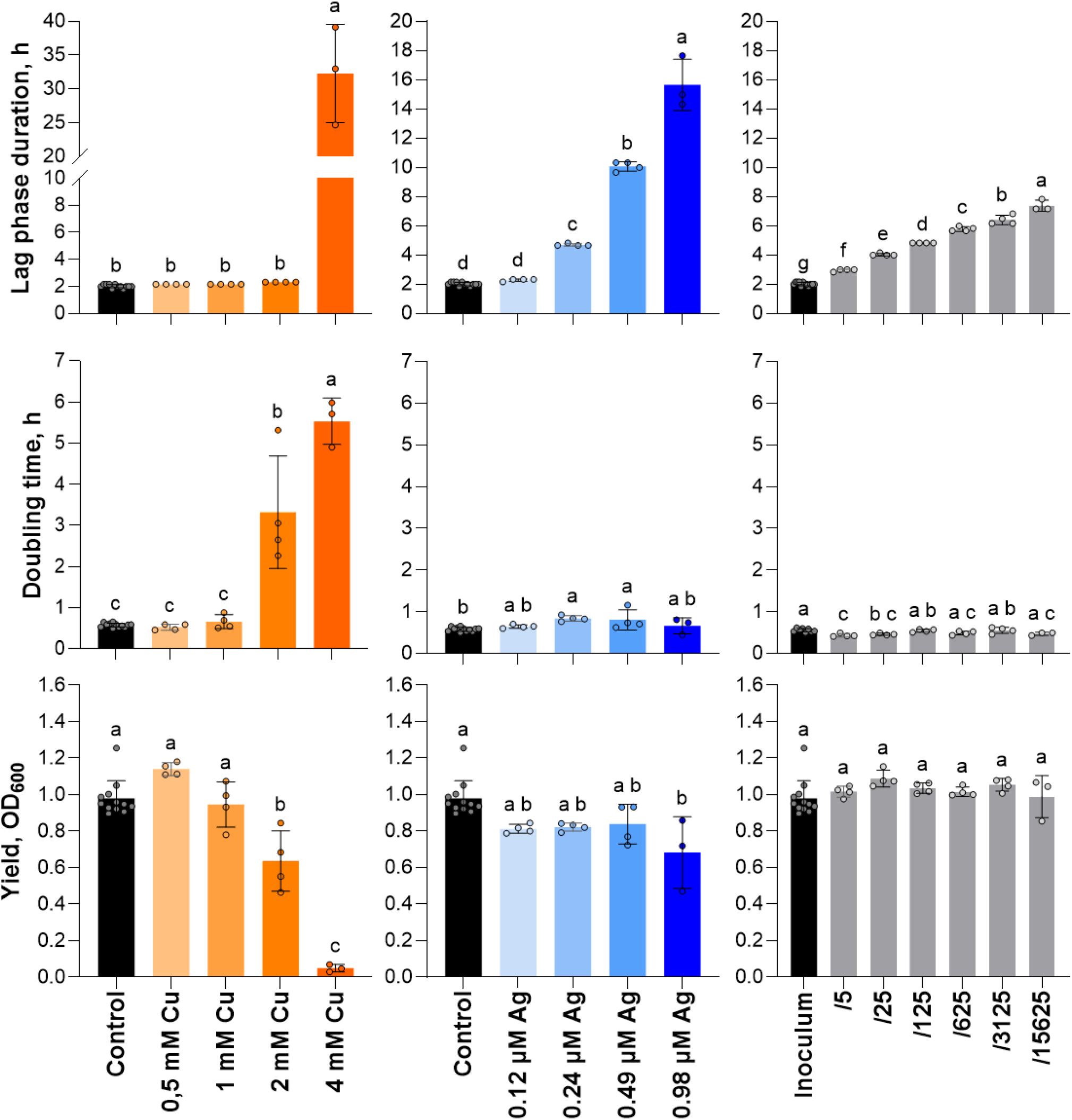
Growth parameters of *E. coli* BW25113 exposed to CuSO_4_ (orange) or AgNO_3_ (blue) or inoculums with different viable cell densities (grey). Values are calculated from growth curves on Supplementary Figures S3 and S4. Nominal metal concentration or inoculum fold dilution is marked on X-axis where appropriate. Single data points with mean and SD are shown. Lower case letters denote similarity groups based on one-way ANOVA followed by Tukey post-hoc test for multiple comparisons at α=0.05.

**Supplementary Figure S2.**
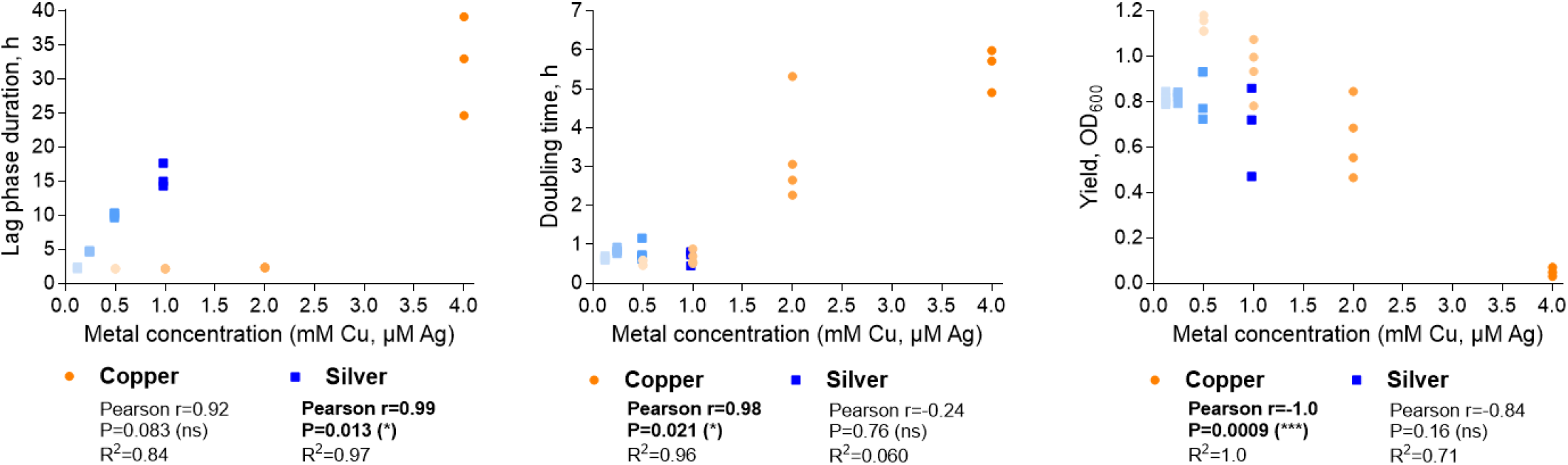
Dose-dependent changes in lag phase duration (left), doubling time (middle) and yield (right) in response to sub-MIC copper (orange gradient) or silver (blue gradient) concentration. Statistically significant linear correlations are marked in bold. Growth parameter values calculated from growth curves on Supplementary Figure S3 and presented on Supplementary Figure S1 were used for correlations. Results from 4 biological replicates are presented.

**Supplementary Figure S3.**
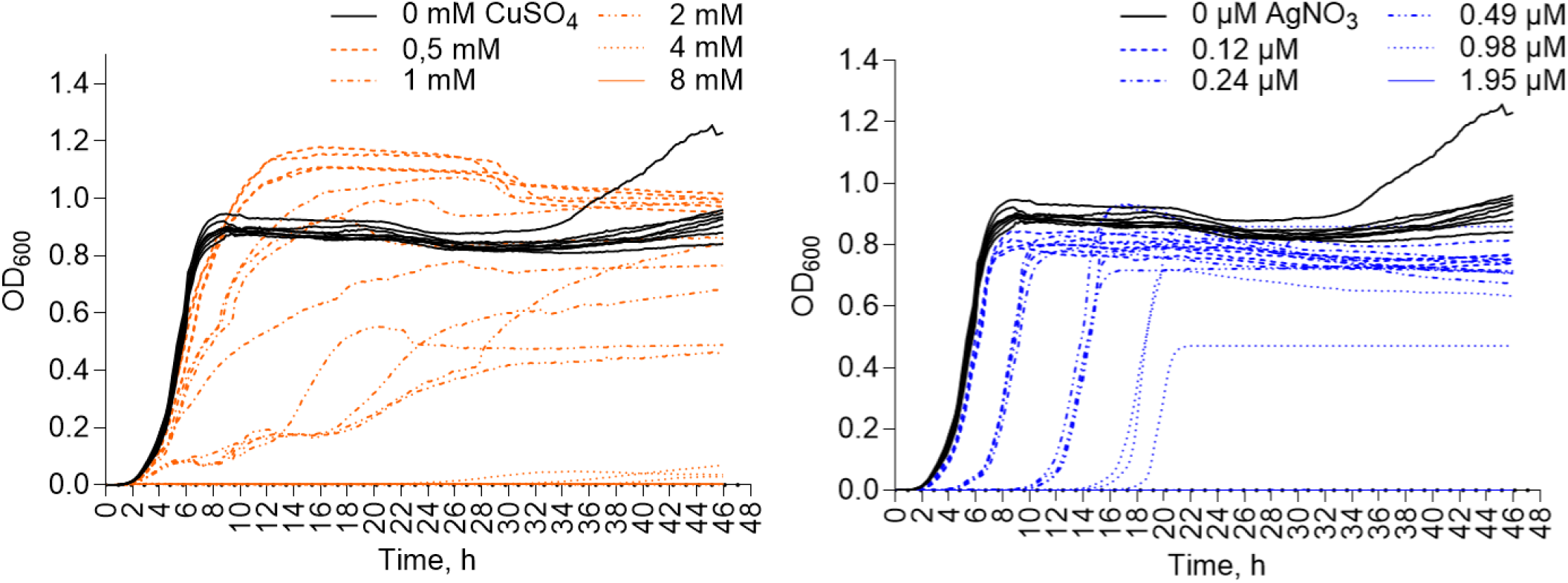
Growth curves of *E. coli* BW25113 in the presence or absence of CuSO_4_ (left, orange) or AgNO_3_ (right, blue). Individual curves from 4 experiments underlying the mean curves on Figure 1 are shown. Non-exposed controls (black) on both panels are pooled across all experiments.

**Supplementary Figure S4.**
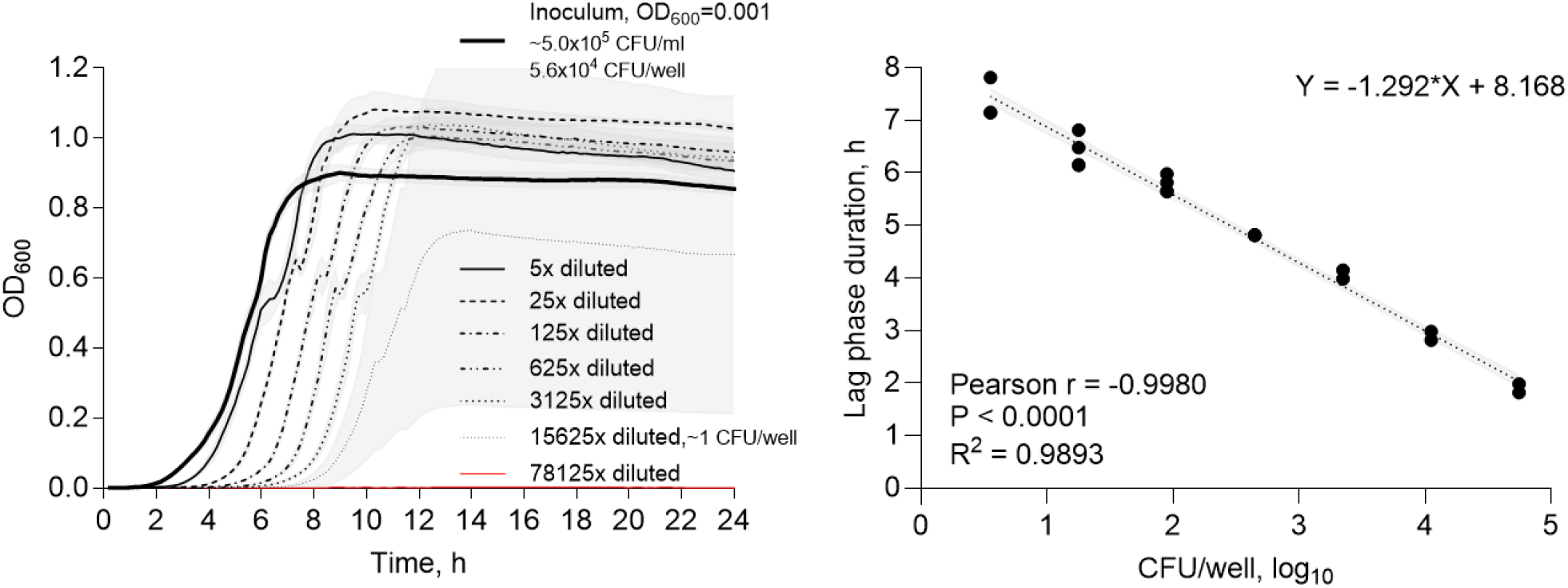
Effect of inoculum density of *E. coli* BW25113 on growth curves (left) and observed strong negative correlation between the lag phase duration and inoculum density (right). Dotted line denotes linear regression. Results from 4 biological replicates are presented.

**Supplementary Figure S5.**
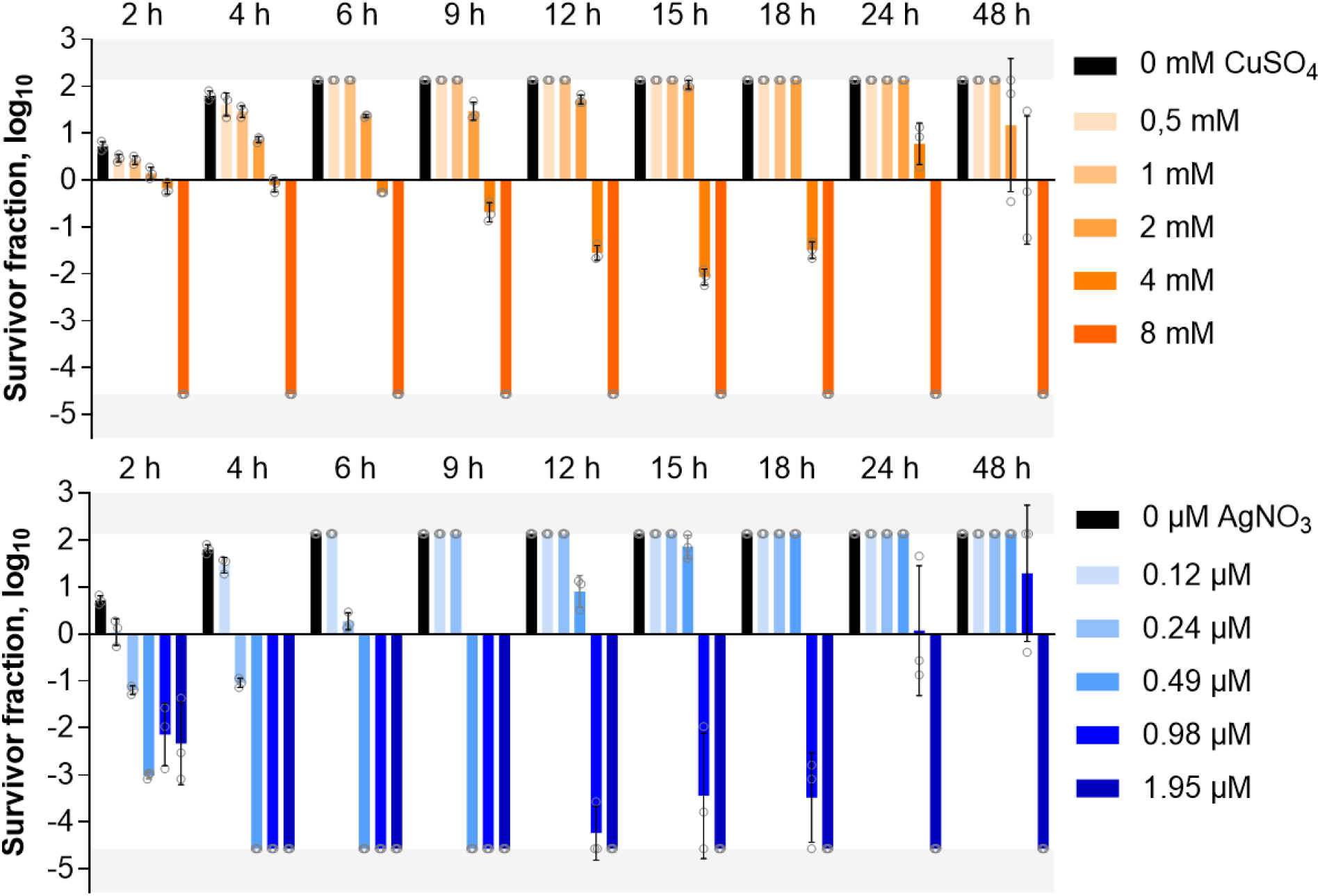
Survival of *Escherichia coli* BW25113 (mean 5.74 log_10_CFU/ml) exposed to CuSO_4_ (orange, upper panel) or AgNO_3_ (blue, lower panel) during 48 h exposure. Mean and SD of 3 experiments underlying the kill curves on Figure 1 are shown. Areas highlighted in grey denote ranges exceeding colony counting detection limits.

